# Tbet promotes NK cell egress from the bone marrow and CXCR6 expression in immature NK cells

**DOI:** 10.1101/583575

**Authors:** Antonia O. Cuff, Thibaut Perchet, Simone Dertschnig, Rachel Golub, Victoria Male

## Abstract

Tbet-deficient mice have reduced NK cells in blood and spleen, but increased NK cells in bone marrow and lymph nodes, a phenotype that is thought to be due to defective migration. Here, we revisit the role of Tbet in NK cell bone marrow egress. We definitively show that the accumulation of NK cells in the bone marrow of Tbet-deficient (Tbx21^-/-^) animals occurs because of a cell-intrinsic migration defect. We identify a profile of gene expression, co-ordinated by Tbet, which affects the localisation of NK cells in the bone marrow. Tbet promotes Cxcr6 expression and immature NK cells accumulate in the bone marrow of CXCR6-deficient mice. This suggests that CXCR6 is among the mediators of migration, controlled by Tbet, that co-ordinate NK cell bone marrow egress.

## Introduction

Tbet was originally described as the key transcription factor directing Th1 lineage commitment (Szabo *et al*, 2000). More recently, it has become clear that Tbet also drives differentiation and memory cell generation in a number of lymphocyte lineages (Kallies and Good-Jacobson, 2017) as well as being required for the development and survival of ILC1 (Spits *et al*, 2013). There is still debate about the extent to which ILC1 form a separate lineage from NK cells (Vivier *et al*, 2018; Gao *et al*, 2017; Cortez *et al*, 2017; Park *et al*, 2019; Xu *et al*, 2019), with one key factor that distinguishes NK cells from ILC1 being the greater extent to which ILC1 depend upon Tbet for their development (Sojka *et al*, 2014; Daussy *et al*, 2014). Nevertheless, Tbet-deficient mice do display defects in NK cell number, maturation status and function (Townsend *et al*, 2004; Jenne *et al*, 2009).

Tbet-deficient mice have reduced NK cells in blood and spleen, but increased NK cells in the bone marrow and lymph nodes (Townsend *et al*, 2004; Jenne *et al*, 2009). These observations led to the suggestion that Tbet is required for NK cells to leave the bone marrow or lymph nodes and enter the blood (Jenne *et al*, 2009). Tbet-deficient NK cells express lower levels of S1pr5 mRNA than their wild type counterparts, and S1pr5 knockout mice phenocopy Tbet-deficient animals, suggesting that Tbet mediates bone marrow and lymph node egress by upregulating S1PR5 expression (Jenne *et al*, 2009). We were prompted to revisit the role of Tbet in bone marrow egress by the unexpected finding that a conditional knockout of Tbet in NK cells did not display the accumulation of NK cells in the bone marrow that has been reported in other Tbet-deficient strains of mice. Here, we report that in the absence of Tbet, NK cells display a cell-intrinsic defect in their ability to leave the bone marrow, and in their ability to differentiate to the final stage of NK cell development. We find that, in the absence of Tbet, CXCR6-expressing bone marrow NK cells are lost. We also observe an accumulation of immature NK cells in the bone marrow in the absence of CXCR6, although this is smaller than that observed in the absence of Tbet, and a reduced ability of CXCR6-deficient bone marrow to reconstitute peripheral NK cell compartments, pointing to a minor role for CXCR6 in NK cell trafficking.

## Results

### Tbet-deficient NK cells accumulate in the bone marrow

NK cells have previously been reported to accumulate in the bone marrow of Tbet-deficient strains of mice (Townsend *et al*, 2004; Jenne *et al*, 2009). We examined the frequencies of Lineage-negative NK1.1^+^ cells within the bone marrow of both Tbx21^-/-^ and Ncr1^iCre^ Tbx21^fl/fl^ mice, compared to appropriate controls (Fig. 1a). Within the lineage-negative NK1.1^+^ compartment, we defined CD11b^-^ cells as immature (iNK) cells, further subdividing the CD11b^+^ mature (mNK) compartment by their expression of CD27 into a CD27^+^ subset of intermediate maturity, which for simplicity is sometimes called “mNK1”, and a more mature CD27^-^ subset called “mNK2” (Kim *et al*, 2002; Chiossone *et al*, 2009; Goh and Huntington, 2017). Eomes^-^ CD49a^+^ cells, suggested to represent ILC1, have previously been reported within the Lineage-negative NK1.1+ bone marrow compartment (Klose *et al*, 2014), although other studies have failed to find a prominent ILC1 population within this gate (Turchinovich *et al*, 2018; Wang *et al*, 2018). In the bone marrow of our mice, we did not find a significant Eomes^-^ CD49a^+^ population (Fig. 1a – the liver, in which a substantial ILC1 population is present, is shown as a positive control). This confirms that, in these animals, this gating strategy identifies NK cells.

**Fig. 1.**
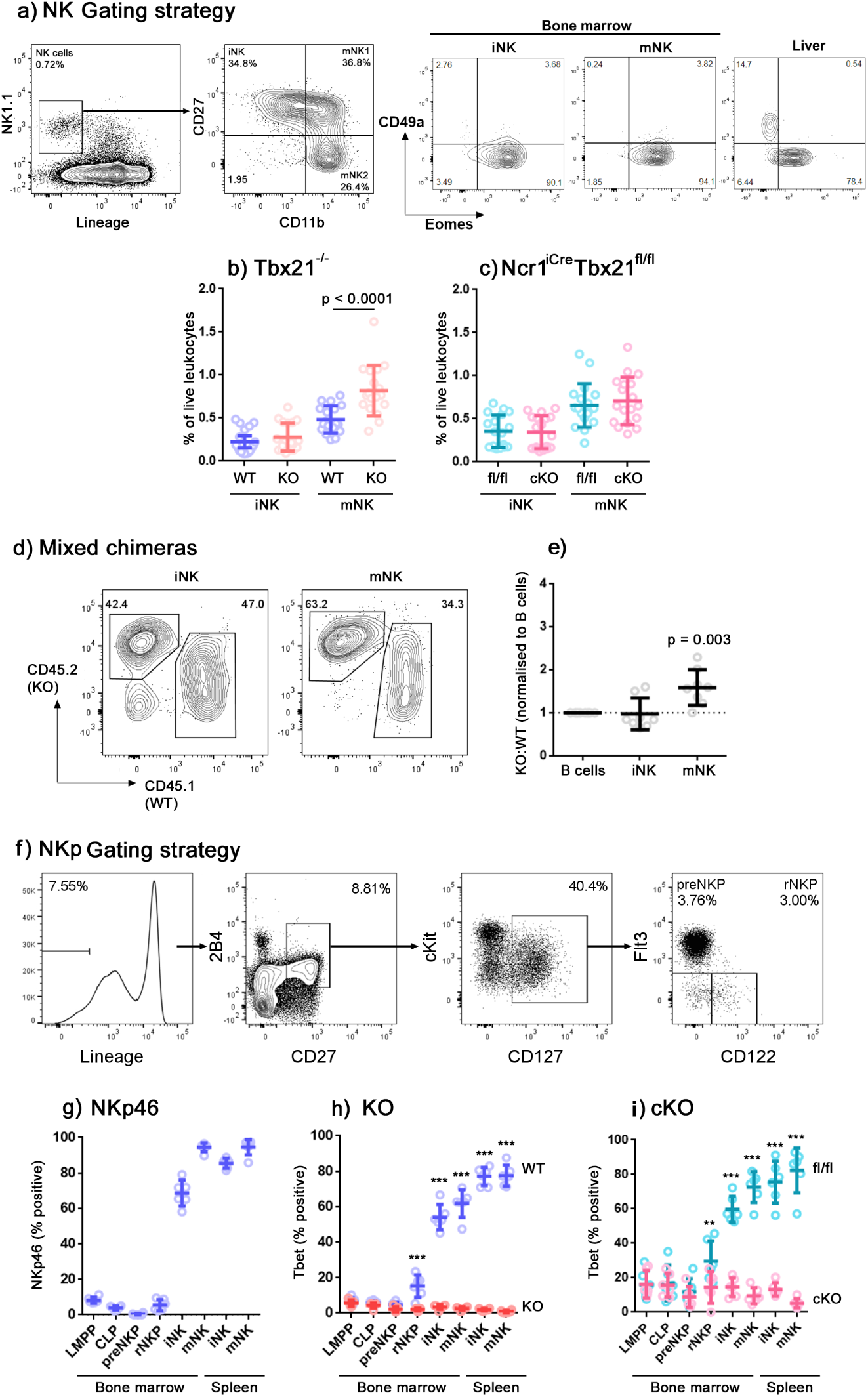
Cell intrinsic accumulation of NK cells in bone marrow in the absence of Tbet. **(a)** Flow cytometry gating strategy identifying NK cells in the bone marrow. NK cells were identified by gating on single, live, CD45^+^ cells and by leukocyte scatter, lineage negative and NK1.1^+^. Constituent subsets iNK, mNK1 and mNK2 were identified by differential CD11b and CD27 expression. Among subsets identified in this way, no significant CD49a^+^ Eomes^-^ ILC1 population was present. Total lineage negative NK1.1^+^ cells in the liver, among which ILC1 are prominent, are shown for comparison. **(b and c)** Summary graphs showing NK cells (iNK and CD11b^+^ mNK) as a percentage of live leukocytes in Tbx21^+/+^ (WT) compared to Tbx21^-/-^ (KO) mice (b) and in Tbx21^fl/fl^ (fl/fl) compared to Ncr1^iCre^ Tbx21^fl/fl^ (cKO) (c). n = 18 mice per group, means and SD are shown. Significance was determined using two sample, one-tailed t tests with Holm-Sidak correction. **(d)** Wild-type mice were lethally-irradiated and reconstituted with a 1:1 mixture of CD45.2 Tbx21^-/-^ (KO) and CD45.1 Tbx21^+/+^ (WT) bone marrow-derived leukocytes to create mixed chimeras. Representative plots gated on iNK and CD11b^+^ mNK at 8 weeks post-transplant are shown. **(e)** Summary data showing the ratio of Tbx21^-/-^ (KO) to Tbx21^+/+^ (WT) in iNK and CD11b^+^ mNK cells, normalised to CD19^+^ B cells according to the method by Jenne *et al*, 2009. n = 8 mice, means and SD are shown. Significance was determined using a single sample, one-tailed t test. **(f)** Gating strategy for identification of NK progenitors (Fathman *et al*, 2011). **(g)** NKp46 expression over development stages from the bone marrow to the peripheral splenic NK cells in wild type mice. **(h and i)** Tbet expression over developmental stages in Tbx21^+/+^ (WT) compared to Tbx21^-/-^ (KO) mice (h), and in Tbx21^fl/fl^ (fl/fl) compared to Ncr1^iCre^ Tbx21^fl/fl^ (cKO) mice (i). (g-i) n = 6, means and SD are shown. Significance was determined using one-tailed t tests with Holm-Sidak correction. * p < 0.05; ** p < 0.01; *** p < 0.001.

In line with previous reports, we found an accumulation of Lineage-negative NK1.1^+^ cells in the bone marrow of Tbx21^-/-^ mice, which predominantly affects the CD11b^+^ mNK population (Fig. 1b). In the bone marrow of Ncr1^iCre^ Tbx21^fl/fl^ conditional knockout mice in which Cre recombinase-mediated excision of Tbx21 (Tbet) is driven by the Ncr1 (NKp46) promoter, we did not observe the accumulation of NK cells that we saw in Tbx21^-/-^ mice (Fig. 1c). The most obvious explanation for this difference would be if the requirement for Tbet in NK cell egress from the bone marrow were not cell-intrinsic. In a previous study, mixed chimera experiments convincingly showed that the requirement for Tbet in NK cells leaving the lymph nodes is cell-intrinsic, but for bone marrow egress the results were less clear (Jenne *et al*, 2009). Therefore, we created mixed chimeras to determine whether the requirement for Tbet in bone marrow egress is cell-intrinsic (Fig. 1d,e). In the chimeras, we saw an increase in the ratio of Tbet-deficient:Tbet-sufficient cells within the CD11b^+^ mNK compartment of the bone marrow, but not within the iNK compartment. This mirrors our observations in Tbx21^-/-^ mice, where an accumulation of NK cells is observed in the mNK, but not in the iNK compartment. These results therefore support the idea that the requirement for Tbet in NK egress from the bone marrow is cell-intrinsic.

In contrast to Tbx21^-/-^ mice, in which Tbet is consistently absent, Tbet depletion in conditional knockout mice is dependent on Ncr1 expression. We examined the expression of NKp46 and Tbet in LMPPs, CLPs and NK progenitors (gating shown in Fig. 1f). NKp46 (encoded by Ncr1) is not detectable at the protein level until the immature NK cell stage of development (Fig. 1g). Therefore we considered the possibility that transient Tbet expression early in development, before Ncr1-mediated Cre recombinase activity is switched on, could be sufficient to allow NK cell egress later in development. A similar situation occurs for the transcription factor Nfil3, whose expression is required before NK-lineage commitment for later development into functional NK cells (Male *et al*, 2014). NK cells in conditional knockouts of Nfil3 under the Ncr1 promoter develop comparably to controls, whereas they fail to develop in mice globally deficient in Nfil3, because Nfil3 is required earlier in NK cell development than it is excised in these mice (Firth *et al*, 2013).

To address whether the requirement for Tbet, like that of Nfil3, occurs at a specific developmental stage, we examined bone marrow leukocytes from Ncr1^iCre^ Tbx21^fl/fl^ mice for Tbet expression at various stages of NK development. We found significant loss of Tbet in NK-lineage committed progenitors, rNKP, in both conditional knockout and knockout mice relative to their controls, and notably before Ncr1 is detectable at the protein level (Fig. 1h,i). Since the deletion of Tbet occurs earlier than might be expected in conditional knockout mice, this indicates that Ncr1, and consequently Cre-recombinase, is transcribed at an earlier developmental stage than the protein reaches a level that can be detected by flow cytometry. This earlier-than-expected loss of Tbet expression also rules out the possibility that the differences in NK cell accumulation that we observe between the global and conditional knockouts are caused by transient expression of Tbet.

### NK cells are less able to leave the bone marrow in the absence of Tbet

Previous reports attribute the bone marrow accumulation of Tbet-deficient NK cells to a defect in migration involving S1PR5, since S1pr5 mRNA is underexpressed in Tbet-deficient NK cells, S1pr5^-/-^ mice phenocopy the accumulation of NK cells in the bone marrow of Tbet-deficient mice and in S1pr5^-/-^ mice the accumulation is caused by a migration defect (Jenne *et al*, 2009; Walzer *et al*, 2007; Mayol *et al*, 2011). However, a defect in NK cell migration from the bone marrow in the absence of Tbet has not yet been formally shown. Therefore, we examined NK cell migration from the bone marrow *in vivo*. Recently-migrated cells translocate to the sinusoids before exiting the bone marrow, so sinusoidal leukocyte staining acts as an indicator of recent migration from the bone marrow. The procedure involves intravenous injection of fluorescently-labelled anti-CD45.2 monoclonal antibody followed by a 2-minute incubation to allow the circulation to carry the antibody to the sinusoids and stain leukocytes in the blood and sinusoids, while sparing parenchymal leukocytes (Fig. 2a) (Pereira *et al*, 2009).

**Fig. 2.**
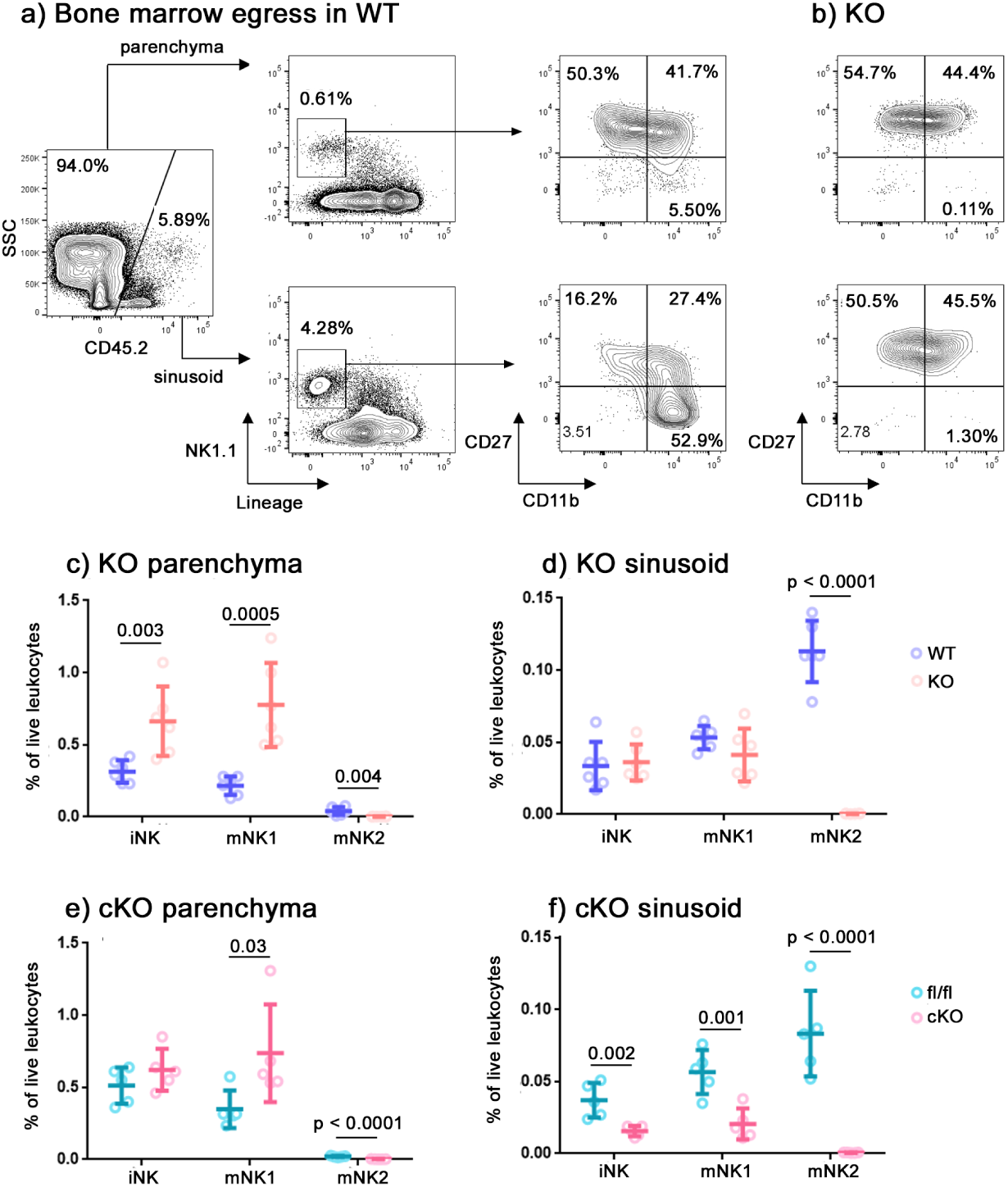
Reduced NK cell bone marrow egress in the absence of Tbet. Sinusoidal NK cells were labelled *in vivo* by intravenous injection of 1µg PE-eFluor610-labelled CD45.2 antibody followed by a 2 minute incubation. **(a)** Representative flow cytometry gating strategy used to identify the NK cells in bone marrow parenchyma (CD45.2-negative) and those which have recently migrated to the sinusoids (CD45.2-positive), a surrogate indicator of egress from the bone marrow in Tbx21^+/+^ (WT) mice. **(b)** NK cells in the parenchyma and sinusoid of Tbx21^-/-^ (KO) mice. **(c - f)** Summary graphs showing the percentage of NK cells as a proportion of live leukocytes in the bone marrow parenchyma (c,e) and sinusoid (d,f) of Tbx21^+/+^ (WT) compared to Tbx21^-/-^ (KO) mice (c,d), and Tbx21^fl/fl^ (fl/fl) compared to Ncr1^iCre^ Tbx21^fl/fl^ (cKO) mice (e,f). n = 6 (c,d) or 5 (e,f) mice per group, mean and SD are shown. Significance was determined using two sample, one-tailed t tests with Holm-Sidak correction.

As NK cells mature from CD11b^-^ iNK to CD11b^+^CD27^+^ mNK1 and then to CD11b^+^CD27^-^ mNK2 in wild type mice, their frequencies decrease in the parenchyma (Fig. 2a,c) and increase in the sinusoids (Fig. 2a,d). This indicates migration out of the bone marrow occurring at all NK cell developmental stages but increasing in more mature cells. We did not observe this pattern in Tbx21^-/-^ NK cells (Fig. 2b), whose frequency instead remained constant between iNK and mNK1 in both parenchyma and sinusoid, indicating a reduced ability to migrate. We also observed a decrease in the frequency of mNK2 in both anatomical compartments, consistent with a defect at the mNK1 to mNK2 transition that has previously been reported (van Helden *et al*, 2015). In the conditional knockout mouse compared to the control, we observed a failure of developing NK cells to leave the parenchyma and enter the sinusoid (Fig. 2e,f), similar to that in the conventional knockout. Therefore, this more sensitive test of bone marrow egress reveals that conditional knockout NK cells, like their conventional knockout counterparts, do have a defect in leaving the BM, although we had been unable to detect this by a simple examination of NK cell frequencies (Fig. 1c).

### Tbet controls the expression of cell migration mediators in bone marrow NK cells

Having confirmed that the accumulation in the bone marrow of Tbx21^-/-^ mice is caused by a migration defect, we sought to define the mediators involved. We therefore sorted iNK and mNK1 cells from the bone marrow of wild type and Tbx21^-/-^ mice, and mNK2 (which are not present in Tbx21^-/-^; Fig. 2a-d) from wild type mice and analyzed them by RNAseq. Raw RNAseq data and differentially expressed gene lists are available from the National Center for Biotechnology Information Gene Expression Omnibus under accession no. GSE122874. Consistent with previous reports (Jenne *et al*, 2009), we found S1pr5 mRNA underexpressed by a factor of approximately seven-fold in both iNK (Fig. 3a,c) and mNK1 (Fig. 3b,c) cells in Tbx21^-/-^ mice compared to wild type. More severely affected by the absence of Tbet was Cxcr6, which was underexpressed by a factor of 125-fold in iNK cells (Fig. 3a,c). The expression of Kit, often considered a marker of immature NK cells, was increased in the absence of Tbet (Fig. 3a,b,d) whereas the expression of genes associated with mature NK effector function, such as Ifng, Gzmb and Prf1, were reduced (Fig. 3a,b,e). Consistent with reports that Tbet promotes Zeb2 (van Helden *et al*, 2015) and antagonizes Eomes (Pikovskaya *et al*, 2016) transcription, Zeb2 and Eomes expression were decreased and increased, respectively, in Tbx21^-/-^ NK cells (Fig. 3a,b,f).

**Fig. 3.**
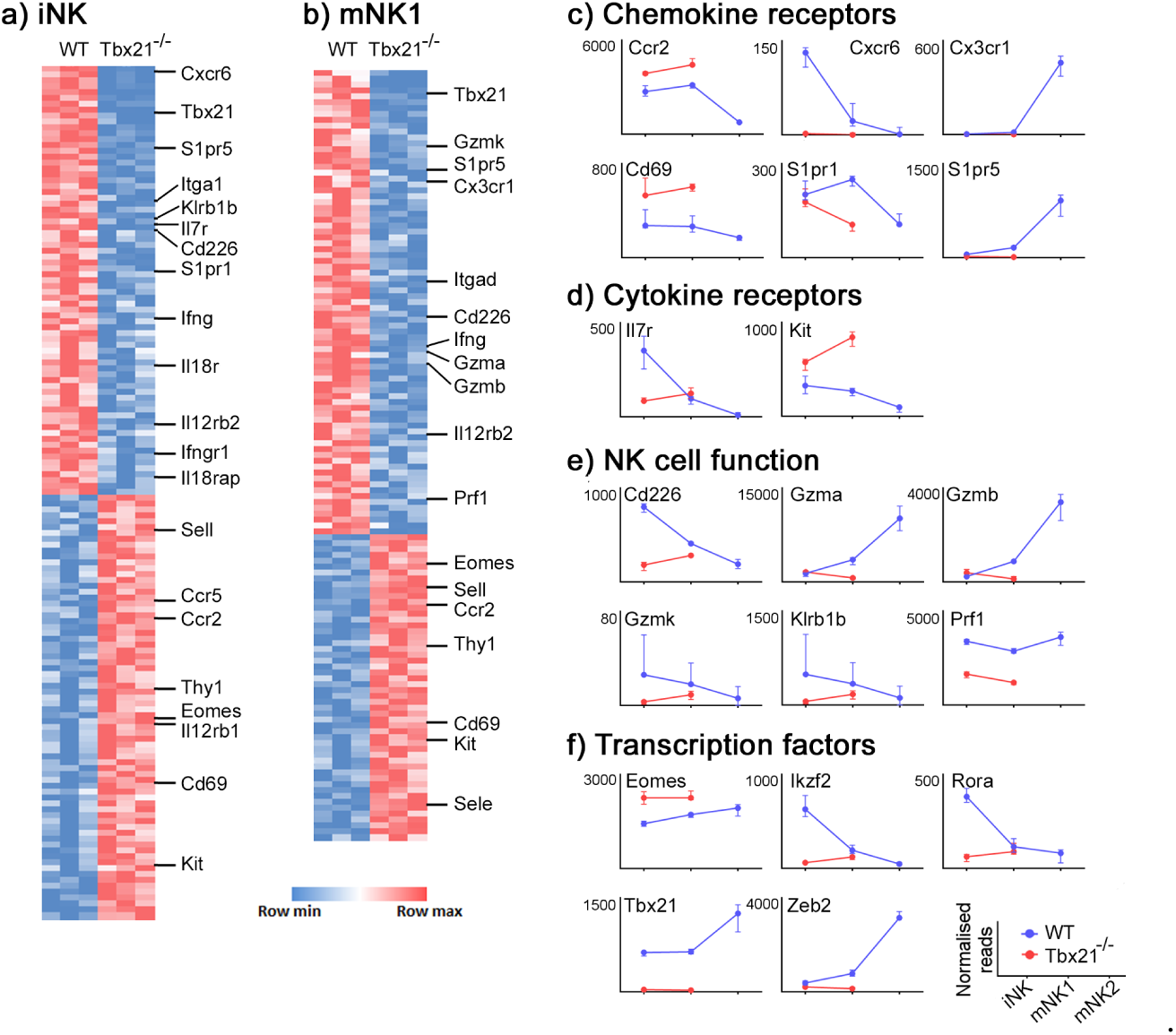
Transcriptional profile of NK developmental stages in the bone marrow of Tbx21^+/+^ and Tbx21^-/-^ mice. **(a**,**b)** RNASeq data from iNK (a) and mNK1 (b) sorted from Tbx21^+/+^ and Tbx21^-/-^ mice. mice (n = 3 pairs). Differentially expressed genes were identified as those with fold change > 2 and p_adj_ < 0.05. **(c-f**) Normalised expression of the transcripts of selected chemokine receptors (c), cytokine receptors (d), NK cell effector proteins (e) and transcription factors (f) over the course of NK cell development are shown (medians and IQR).

We next selected genes identified as differing transcriptionally between wild type and Tbx21^-/-^ NK cells to confirm the differences in their expression at the protein level by flow cytometry (Fig. 4). The genes were selected based on a significance cutoff of p_adj_ < 0.05 and a two-fold or more differential expression between wild type and Tbx21^-/-^ iNK and mNK1 cells. Sell and Cd69 did not have a two-fold difference between KO and WT cells, but were included in the analysis because we observed differential transcription of these genes in both NK subsets.

**Fig. 4.**
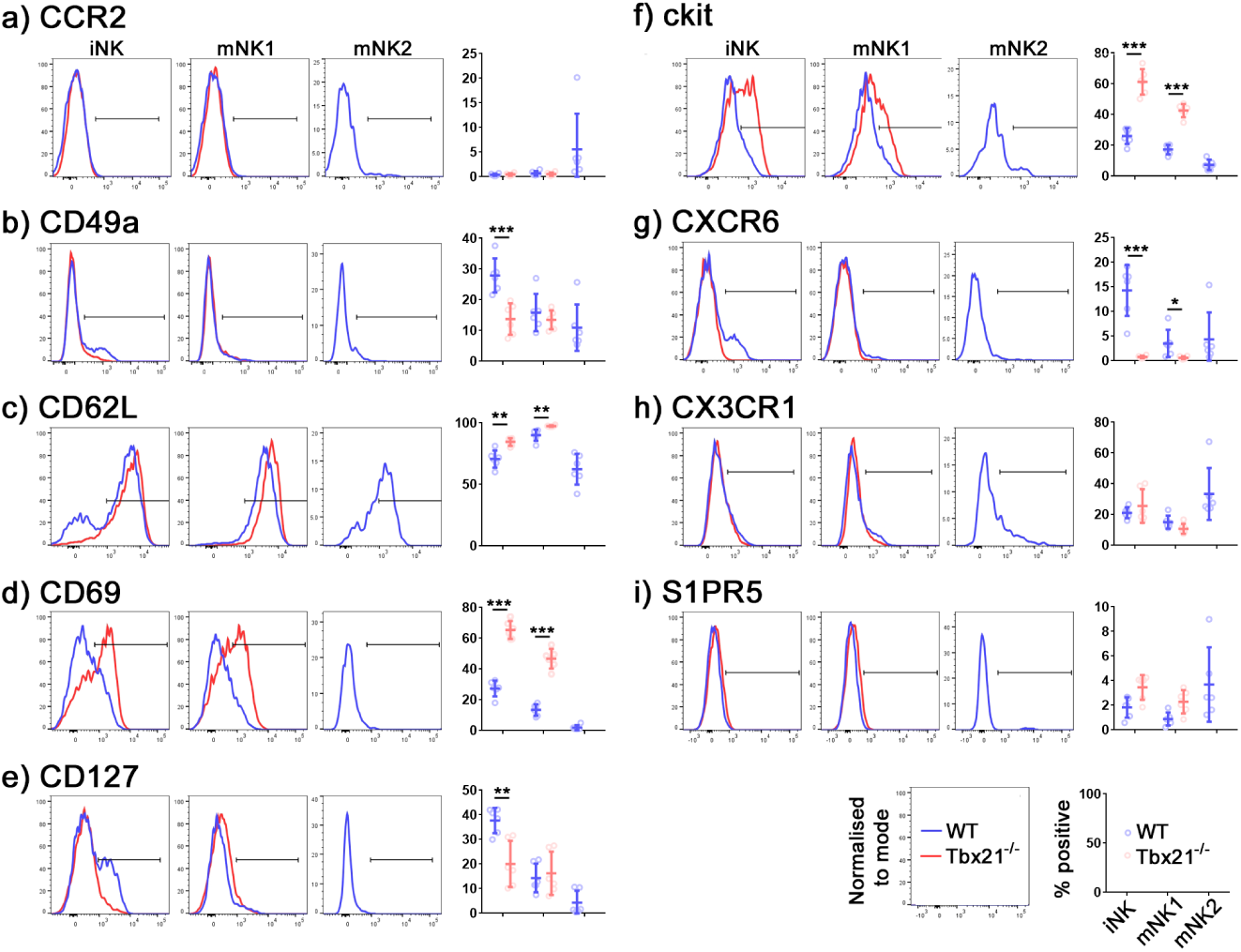
Phenotype of developing NK cells in the presence or absence of Tbet. Leukocytes were isolated from Tbx21^+/+^ (WT; blue) and Tbx21^-/-^ (red) bone marrow and iNK, mNK1 and mNK2 cell subsets were screened by flow cytometry for differentially expressed proteins. Proteins examined were selected based on differential expression of their transcripts between Tbx21^+/+^ and Tbx21^-/-^ in the RNASeq screen. Representative histograms and summary graphs for **(a)** CCR2, **(b)** CD49a, **(c)** CD62L, **(d)** CD69, **(e)** CD127, **(f)** cKit, **(g)** CXCR6, **(h)** CX3CR1, and **(i)** S1PR5 are shown. n = 6 mice per group, means and SD are shown. Significance was determined using two-sample, one-tailed t tests with Holm-Sidak correction. * p < 0.05, ** p < 0.01, *** p < 0.001.

The expression of several proteins associated with cell migration differed between wild type and knockout NK cells. In particular, CXCR6 was expressed at a lower level by knock-out iNK and mNK1, compared to their wild type counterparts (Fig. 4g). Consistent with the RNASeq data, the most significant drop in expression was within the iNK cell compartment. CD69, which suppresses the expression of S1PR1 in order to retain immune cells in lymph nodes and tissues, is upregulated in both knockout NK subsets (Fig. 4d). CD62L (encoded by Sell), which is involved in leukocyte homing from the blood to tissues, also has elevated expression in knockout NK cells compared to wild type (Fig. 4c). We also observed a decrease in the expression of CD49a in knockout iNK (Fig. 4b).

S1pr5 and Ccr2 displayed relatively large differences at the RNA level (Fig. 3a,b), but using antibodies validated on appropriate positive control cells (S1pr5-transfected 293T cells and CD11b^+^ Ly6G^-^ monocytes, respectively) we did not see any differences at the protein level (Fig. 4a,i). Neither did we see any significant difference in CX3CR1 expression between wild type and knockout cells, although the expression of this protein did increase in mNK2 compared to previous stages of development (Fig. 4h).

### Immature NK cells accumulate in the bone marrow of CXCR6-deficient mice

We have previously reported that ILC progenitors require CXCR6 to leave the bone marrow (Chea *et al*, 2015), so we were intrigued by the underexpression of CXCR6 in Tbx21^-/-^ NK cells, which are also less able to leave the bone marrow. If CXCR6 is one of the mediators through which Tbet controls NK cell bone marrow egress, we would expect to see an accumulation of NK cells in the bone marrow of CXCR6-deficient mice, similar to our previous observations of ILC progenitors.

To test this hypothesis, we examined NK cells in the bone marrow of Cxcr6^gfp/gfp^ mice, in which both Cxcr6 alleles are inactivated and replaced by a reporter cassette encoding GFP, or Cxcr6^gfp/+^ controls. We did not observe an accumulation among the general population of CLP, αLP1 or αLP2 in the bone marrow of these mice, although we did observe an accumulation specifically among the Cxcr6-gfp^+^ population of αLP2 (Fig. 5a). Examining NK cell progenitors, we observed an accumulation of iNK, although this was not significant. However, we did observe a significant accumulation specifically among the Cxcr6-gfp^+^ population of iNK (Fig. 5b). In contrast to the Tbet knockout, we did not detect an accumulation of cells among either of the mature NK cell subpopulations in the bone marrow of CXCR6-deficient mice, suggesting that any requirement for CXCR6 in bone marrow egress occurs relatively early in NK cell development.

**Fig. 5.**
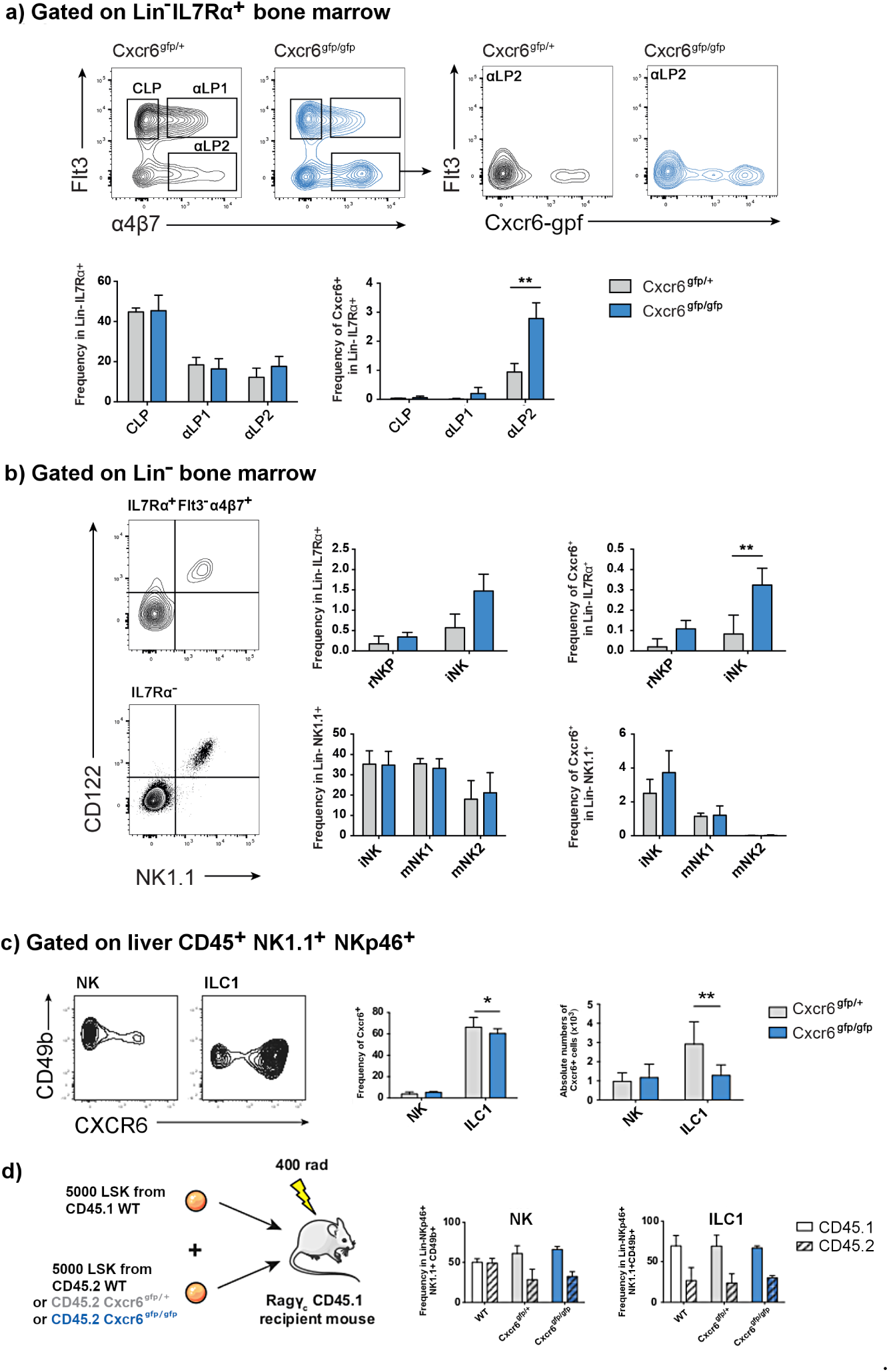
Accumulation of Cxcr6^+^ iNK in the bone marrow of CXCR6-deficient mice. **(a)** Representative flow cytometry gating strategy used to identify CLP, αLP1, αLP2 progenitors (Lin^-^ IL7Rα^+^) in the Cxcr6^gfp/+^ and Cxcr6^gfp/gfp^ bone marrow (Yu *et al*, 2014). αLP2 cells were identified as Flt3^-^α4β7^+^ cells and contour plots indicate the fraction of Cxcr6-gfp^+^ cells among αLP2 from both Cxcr6^gfp/+^ and Cxcr6^gfp/gfp^ mice. Frequencies of the CLP, αLP1 and αLP2 are not significantly different among total bone marrow progenitors. When progenitors are selected as Cxcr6-expressing cells, accumulation of αLP2 fraction in the bone marrow is significant. n = 10 mice per group, means and SD are shown. **(b)** NK progenitor subsets among Lin^-^ IL7Rα^+^ or the Lin^-^ IL7Rα^-^ subsets are identified using CD122 and NK1.1 expression as shown. rNKP and iNK are identified among the IL7Rα^+^ Flt3^-^α4β7^+^ fraction as CD122^+^NK1.1^-^ and CD122^+^NK1.1^+^ subsets, respectively. Among IL7Rα^-^ cells, iNK, mNK1 and mNK2 are identified as shown in Figure 1. In CXCR6-deficient mice, iNK frequency presents a tendency to increase with a significant increase for the Cxcr6-gfp^+^ IL7Rα^+^ iNK fraction. n = 10 mice per group, means and SD are shown. Significance was determined using two sample, one-tailed t tests. * p < 0.05, ** p < 0.01, *** p < 0.001. **(c)** CXCR6 expression in liver NK cells and ILC1. Frequency as a percentage of parent population (CD45^+^ NK1.1^+^ NKp46^+^ CD49b^+^ for NK cells; CD45^+^ NK1.1^+^ NKp46^+^ CD49b^-^ for ILC1) and absolute number of Cxcr6-gfp-expressing ILC1 are significantly decreased in the liver of CXCR6-deficient mice. n = 20 mice per group, means and SD are shown. **(d)** CXCR6 deficiency impacts hepatic NK cell reconstitution. Schematic representation of experimental procedure for competitive reconstitution of congenic lymphopenic Rag/γc double KO mice. Frequencies of CD45.1 (WT reconstitution control) and CD45.2 (WT, Cxcr6^gfp/+^ or Cxcr6^gfp/gfp^ experimental) cells among reconstituted hepatic NK cells and ILC1 are shown.

CXCR6 is highly expressed by ILC1 (Fig. 5c) and we have previously shown that it is required for reconstitution of ILC3 progenitors (Chea *et al*, 2016). Therefore, we investigated the relative contributions of CXCR6 to reconstitution of ILC1 compared to NK cells. For these experiments, we focused on the liver, since a large ILC1 population is present here. At steady state, CXCR6-deficient mice displayed a defect in ILC1 frequency and number (Fig. 5c), consistent with the requirement for CXCR6 in ILC progenitor trafficking during fetal life that we have previously reported (Chea *et al*, 2016). To determine the requirement for CXCR6 in ILC1 and NK cell reconstitution in adult life, we performed competitive reconstitution experiments (Fig. 5d). Surprisingly, we did not observe any difference in the relative reconstitution of ILC1 in competitive reconstitutions using either CXCR6-sufficient or -deficient progenitors. This suggests that, in contrast to the situation in fetal life, ILC1 in adults do not seem to be dependent on CXCR6 for trafficking. On the other hand, the reconstitution of NK cells was reduced when mice were reconstituted with progenitors that lacked either one or both alleles of Cxcr6, compared to wild type. This points to a role for CXCR6 in NK cell circulation during adult life, with the deficiency in reconstitution observed in heterozygous progenitors suggesting that a threshold of CXCR6 protein expression, higher than that which can be expressed by one allele, may be required to mediate reconstitution.

## Discussion

In this study, we revisited the role of Tbet in mediating NK cell egress from the bone marrow during development. We observed an accumulation of NK cells in the bone marrow of Tbx21^-/-^ mice, in agreement with previous reports (Townsend *et al*, 2004; Jenne *et al*, 2009). The accumulating NK cells in Tbet-deficient mice have previously been described as CD27^hi^ and KLRG1^lo^ (Jenne *et al*, 2009), but no distinction was made between the two CD27^+^ subsets, CD27^+^CD11b^-^ iNK and CD27^+^CD11b^+^ mNK1. Here we report that the accumulation occurs specifically within the mNK1 compartment, although this is unsurprising given a more recent report, which we confirm here, that there is a block at the mNK1 to mNK2 transition in the absence of Tbet (van Helden *et al*, 2015). We confirmed that the requirement for Tbet in NK cell bone marrow egress is cell-intrinsic, demonstrating accumulation of Tbx21^-/-^ NK cells in the bone marrow of mixed bone marrow chimeras more convincingly than in previous studies (Jenne *et al*, 2009). Potentially, this could be because the earlier studies were carried out using a Tbet-deficient strain called Duane, in which Tbet protein levels were reduced by approximately four-fold, whereas Tbx21^-/-^ NK cells completely lack Tbet.

The accumulation of NK cells in the bone marrow of Tbet-deficient mice, together with the cells’ underexpression of S1pr5 mRNA and the defect in bone marrow egress displayed by S1pr5^-/-^ NK cells led to the suggestion that Tbet mediates NK cell bone marrow egress via S1PR5 (Jenne *et al*, 2009; Walzer *et al*, 2007; Mayol *et al*, 2001). However, no previous study had formally addressed the question of whether the accumulation of NK cells in the absence of Tbet occurs due to a migration defect. We used *in vivo* staining of sinusoidal NK cells to show that Tbet-deficient iNK and mNK1 cells are less able to move from the parenchyma to the sinusoid of the bone marrow, supporting the idea that Tbet promotes NK cell bone marrow egress. The *in vivo* bone marrow egress staining further revealed that Tbet knockout iNK cells and conditional knockout iNK and mNK1 cells all have a migration defect that was not apparent on simple examination of cell frequencies at steady state, suggesting that this test detects migration defects with greater sensitivity. We also noted a slight increase in the frequency of NK cells in the parenchyma of Tbx21^fl/fl^ mice, compared to wild type, although the experiments were not designed such that these two could be statistically compared. This slight accumulation of parenchymal NK cells in the control animals may have contributed to the reduced difference between the conditional knockouts and controls on examination of cell frequencies at steady state, despite the defect in NK cell migration still being present.

After we confirmed that the NK cell accumulation in Tbet-deficient bone marrow was caused by defective migration, we explored which mediators were involved, identifying a transcriptional signature associated with Tbet expression in developing NK cells. We confirmed previous reports that S1pr5 is reduced in Tbet-deficient NK cells at the transcriptional level (Jenne *et al*, 2009), but we were unable to corroborate this at the protein level. We were also surprised not to see significant differences in CCR2 and CX3CR1 protein expression, since both differed at the transcript level and both CCR2-deficient and CX3CR1-deficient mice display an accumulation of NK cells in their bone marrow (Ponzetta *et al*, 2013; Fujimara *et al*, 2015). On the other hand, a number of molecules involved in cell migration did differ between wild type and Tbx21^-/-^ NK cells, with increased expression of CD69 and CD62L, which would be expected to retain cells in the parenchyma, as we observed (Cibrián and Sánchez-Madrid, 2017) and decreased expression of CD49a and CXCR6.

We have previously shown that CXCR6 is important for ILC progenitors leaving the bone marrow (Chea *et al*, 2016) so we went on to investigate the role of CXCR6 in NK cell bone marrow egress. In the bone marrow of CXCR6-deficient mice, we found an accumulation of iNK, although this was only significant among Cxrc6-gfp^+^ cells. This accumulation among iNK was consistent with our observation that CXCR6 expression was both highest and most differentially expressed within this population. We did not observe an accumulation among the CD11b^+^ mNK populations. Therefore, the NK cell accumulation we observed in the absence of CXCR6 was less pronounced than that in the absence of Tbet, and was only observed in early developmental stages. This suggests that although CXCR6 may be one of the molecules through which Tbet mediates NK cell bone marrow egress, its role is likely to be relatively minor, with other targets of Tbet, such as S1pr5 and Cd69, playing a greater role at later stages of NK cell development.

One key question which could affect the interpretation of these results is the extent to which the CXCR6^+^ cells which we found within the iNK subset could represent Tbet-dependent ILC1, significant numbers of which have been reported by some studies (Klose *et al*, 2014), although others have found smaller or negligible numbers (Turchinovich *et al*, 2018; Wang *et al*, 2018). The variation in these reports may result from different ages of animals being used in the experiments, since ILC1 are more dominant in younger animals (Constantinides *et al*, 2015), or even from as-yet-undefined colony-specific factors. In the bone marrow of the animals used in this study, we were unable to find a significant Eomes^-^ CD49a^+^ ILC1 population. Indeed, the CD49a^+^ cells were identified in this study were also positive for CD49b, similar to a previous report, which found these double-positive cells to account for approximately 10% of NK cells in the bone marrow (Wang *et al*, 2018). This supports the idea that the cells we are characterizing in the bone marrow in this study are, indeed, NK cells. Our observations of CXCR6-deficient mice further support the idea that this is an NK cell and not an ILC1 phenomenon, since in adult life CXCR6-deficient bone marrow progenitors are somewhat less able to reconstitute the NK cell compartment than their CXCR6-sufficient counterparts, whereas they are equally able to reconstitute the adult ILC1 compartment. Indeed if, as some recent studies have suggested, NK cells and ILC1 do not form separate lineages, it will not be possible to define phenomena as pertaining to NK cells separately from ILC1 (Gao *et al*, 2017; Cortez *et al*, 2017, Park *et al*, 2019; Xu *et al*, 2019).

Overall, we have shown that Tbet is required for NK cells to leave the bone marrow in a cell-intrinsic manner. We further define a profile of gene expression coordinated by Tbet which promotes NK cell bone marrow egress. Among this group of genes which mediate localization of NK cells within the bone marrow, is Cxcr6. Bone marrow egress of iNK cells is somewhat defective in the absence of CXCR6, but the phenotype occurs both at an earlier developmental stage and is less pronounced that than in the absence of Tbet, suggesting that other targets of Tbet are likely to have a more significant role in the more mature NK cells that are the primary population leaving the bone marrow. As we move into an era where the cancer-fighting potential of NK cells is being harnessed to develop immunotherapies, it will become increasingly important to understand NK egress in order to maximize the full potential of those therapies (Souza-Fonseca-Guimaraes *et al*, 2019) and it will be interesting in the future to better define the roles of some of the other genes that we identified in NK cell bone marrow egress.

## Materials and Methods

### Mice

B6.129S6-Tbx21^<tm1Glm>^/J (RRID IMSR_JAX:004648; “Tbx21^-/-^”) and B6.129-Tbx21^tm2Srnr^/J mice (RRID IMSR_JAX:022741; “Tbx21^fl/fl^”) were purchased from the Jackson Laboratory. B6(Cg)-Ncr1^tm1.1(icre)Viv/Orl^ mice (Narni-Mancinelli *et al*, 2011) (RRID MGI:5309017; “Ncr1^iCre^”) were acquired from the European Mutant Mouse Archive as frozen embryos and rederived at the Royal Free Hospital, London. Ncr1^iCre^ mice were crossed onto Tbx21^fl/fl^ to produce Ncr1^iCre^ Tbx21^fl/fl^ conditional knockouts and Ncr1^WT^ Tbx21^fl/fl^ littermate controls, as previously described [22]. Tbx21^-/-^ mice were crossed onto C57BL/6J mice, bred at the Royal Free Hospital, and the resultant heterozygotes were crossed to produce Tbx21^-/-^ homozygote knockouts and Tbx21^+/+^ homozygote wild type littermate controls. Mice were sacrificed between 6 and 12 weeks of age with direct cervical dislocation. Death was confirmed by cessation of circulation. The hind leg bones (tibia and femurs) and spleen were dissected to isolate leukocytes. Animal husbandry and experimental procedures were performed according to UK Home Office regulations and institute guidelines under project licence 70/8530.

Cxcr6^gfp/+^ and Cxcr6^gfp/gfp^ mice (Geissman *et al*, 2005) were bred in the animal facilities at Pasteur Institute, Paris. Mice were bred in accordance with Pasteur Institute guidelines, in compliances with European welfare regulations, and all animal studies were approved by Pasteur Institute Safety Committee in accordance with French and European guidelines. Mice were sacrificed between 8 and 12 weeks of age with carbon dioxide pump. Death was confirmed by cessation of circulation. The hind leg bones (tibia and femurs) were dissected to isolate leukocytes.

### Bone marrow transplantation

Recipient mice were given myeloablative irradiation before syngeneic bone marrow transplantation at 10 weeks of age. Mice received a total of 11Gy lethal irradiation, 5.5 Gy each on day −2 and day 0. On day 0, mice were reconstituted 4 hrs post-irradiation with 5 million donor bone marrow cells in 200 μl Hank’s Balanced Salt Solution (HBSS; Lonza, distributed by VWR, Lutterworth, UK), administered by intravenous injection into the tail vein. C57BL/6J (“CD45.2”) wild type mice were used as recipient mice, which received a 1:1 mixture of donor bone marrow cells from CD45.1 WT and (CD45.2) Tbx21^-/-^ mice to produce mixed chimeras. Bone marrow transplanted mice were sacrificed and analysed 8 weeks post-transplant.

### Sinusoidal cell labelling

Mice received an intravenous injection of 1 μg anti-CD45.2 PE-eFluor 610 monoclonal antibody (eBioscience) in phosphate buffered saline (PBS; Life Technologies brand, Thermo Fisher Scientific, Waltham, MA, USA). Mice were sacrificed after two minutes.

### Cell isolation

Bone marrow was flushed from dissected bones with supplemented RPMI 1640 medium (Life Technologies). Tissue clumps were pelleted at 500 xg, 4°C for 5 minutes, then mechanically disaggregated and filtered through a 70 μm cell strainer. RPMI 1640 medium was supplemented with 10% fetal calf serum, non-essential amino acids, sodium pyruvate, penicillin-streptomycin, 25 mM HEPES buffer and 50 μM 2-Mercaptoethanol (all from Life Technologies).

For cell sorting on a FACSAria (BD Biosciences, Oxford, UK), isolated bone marrow cells were ACK lysed by incubation in ACK lysing buffer (Life Technologies) for 5 minutes at room temperature to remove excess red blood cells and to enrich for leukocytes before cell sorting. The leukocytes were further enriched for NK cells by immuno-magnetic depletion of lineage-associated cells (CD3, CD8α, CD19, Gr-1) using the relevant FITC-conjugated antibodies and anti-FITC MicroBeads (Miltenyi, Woking, UK). The sorting buffer consisted of PBS, 0.5% bovine serum albumin (BSA; Sigma Aldrich, Hammerhill, UK) and 2 mM EDTA (Sigma Aldrich).

Spleens were passed through a 40 μm cell strainer. Red blood cells were lysed by incubation in ACK lysing buffer for 5 minutes at room temperature.

### Flow cytometry

The following anti-mouse antibodies were used: anti-CCR2-PE-Cyanine7 (clone SA203G11, Biolegend, London, UK), anti-CD3-FITC (17A2, Biolegend), anti-CD3-biotin (clone 145-2C11, Sony Biotechnology, Surrey, UK), anti-CD4-BV786 (clone RM4-5, BD Horizon), anti-CD8α-FITC (clone 53-6.7, Biolegend), anti-CD8α-biotin (clone 53-6, Sony), anti-CD11b-APC (clone M1/70, Biolegend), anti-CD11b-PE (clone M1/70, Biolegend), anti-CD11b-AlexaFluor700 (clone M1/70, BD Pharmingen), anti-CD19-FITC (clone 6D5, Biolegend), anti-CD19-PerCP-Cyanine5.5 (clone 6D5, Biolegend), anti-CD19-biotin (clone 6D5, Sony), anti-CD27-APC (clone LG.3A10, Biolegend), anti-CD27-APCeFluor780 (clone LG.7F9, eBioscience, San Diego, CA), anti-CD27-PE-Dazzle594 (clone LG.3A10, Biolegend), anti-CD27-APC-Cyanine7 (clone LG.7F9, eBioscience), anti-CD45-BV510 (clone 30-F11, Biolegend), anti-CD45-APC-Cyanine7 (clone 30-F11, BD Pharmingen), anti-CD45.2 PE-eFluor610 (clone 104, eBioscience), anti-CD45.1-PE-Cyanine7 (clone A20, Biolegend), anti-CD49a-BUV395 (clone Ha31/8, BD Bioscience), anti-CD49b-BV510 (clone HMα2, BD Optibuild), anti-CD62L-PerCP-Cyanine5.5 (clone MEL-14, Biolegend), anti-CD69-PerCP-Cyanine5.5 (clone H1.2F3, Biolegend), anti-CD117(cKit)-BV510 (clone 2B8, Biolegend), anti-CD117 (ckit)-APC-Cyanine7 (clone ACK2, Sony), anti-CD122 (IL-2Rβ)-eFluor450 (clone TM-β1, eBioscience), anti-CD122 (IL-2R-β)-APC (clone TM-β1, Biolegend), anti-CD127 (IL-7Rα)-PE (clone A7R34, eBioscience), anti-CD127 (IL-7Rα)-PE-Cyanine7 (clone A7R34, Sony), anti-CD135 (Flt3)-PerCP-eFluor710 (clone A2F10, eBioscience), anti-CD135 (Flt3)-PE (clone A2F10, Sony), anti-CD244.2-PE-Cyanine7 (clone m2B4 (B6)458.1, Biolegend), anti-CX3CR1-BV510 (clone SA011F11, Biolegend), anti-CXCR6-PE-Cyanine7 (clone SA051D1, Biolegend), anti-Gr-1-FITC (clone RB6-8C5, Biolegend), anti-Gr-1-APC (clone RB6-8C5, Biolegend), anti-GR1-biotin (clone RB6-8C5, BD Pharmingen), anti-integrin α4β7-BV421 (clone DATK32, BD Horizon), anti-NK1.1-FITC (clone PK136, Biolegend), anti-NK1.1-APC-eFluor780 (clone PK136, eBioscience), anti-NK1.1-BV650 (clone PK136, Sony), anti-NKp46-BV711 (clone 29A1.4, Biolegend), anti-NKp46-V450 (clone 29A1.4, BD Horizon), anti-Sca-1-AlexaFluor700 (clone D7, Biolegend), anti-Sca-1-BV510 (clone D7, Sony), anti-S1PR5-PE (clone 1196A, Bio-Techne, Abingdon, UK), anti-TCRαβ-biotin (clone H57 597, Sony), anti-TCRγδ-biotin (clone GL3, BD Pharmingen), anti-TER119-biotin (clone TER-119, Sony) and streptavidin-PE-Cyanine5 (BD Pharmingen) for surface antigens; and anti-Eomes-PE-Cyanine7 (clone Dan11mag, eBioscience) and anti-Tbet-eFluor660 (clone eBio4B10, eBioscience) for intracellular staining.

In Figures 1-4, the lineage cocktail used to identify iNK, mNK1 and mNK2 consisted of anti-CD3-, CD8α-, CD19- and Gr-1-FITC (Lin1). The lineage cocktail used for the identification of LMPP, CLP, preNKP and rNKP in Figure 1 consisted of anti-CD3-, CD8α-, CD19-, Gr-1- and NK1.1-FITC. Lineage cocktail for Figure 5 consisted of anti-CD3-, CD8α-, CD19-, TCRαβ-, TCRγδ-, GR1- and TER119-biotin, stained with streptavidin-PE-Cyanine5. Dead cells were excluded using fixable viability dye eFluor 450 (eBioscience), fixable viability dye eFluor 455 ultraviolet (eBioscience), Propidium iodide (Sigma Aldrich) or DAPI (4’,6-Diamidino-2-Phenylindole, Dilactate; Biolegend). Intracellular staining was carried out using Human FoxP3 Buffer (BD Biosciences) according to the manufacturer’s instructions. Data were acquired on an LSRFortessa II (BD Biosciences) and analyzed using FlowJo (Tree Star, Ashland, OR). Cells were sorted on a FACSAria (BD Biosciences).

### RNA sequencing

Total RNA was extracted from sorted iNK, mNK1 and mNK2 bone marrow NK cells using an RNeasy Micro kit (Qiagen, Manchester, UK). cDNA Libraries were prepared from 2ng of total RNA (RIN 3.2-10) using the Ovation Solo assay (NuGEN, AC Leek, The Netherlands) with 15 cycles of amplification. Libraries were assessed for correct size distribution on the Agilent 2200 TapeStation and quantified by Qubit DNA High Sensitivity assay (Thermo Fisher Scientific) before being pooled at an equimolar concentration. Sequencing was performed on the Illumina NextSeq 500, generating approximately 12million 75bp single end reads per sample. Differential expression analysis was carried out using SARTools (Varet *et al*, 2016) filtering at padj < 0.05.

## ACKNOWLEDGEMENTS

This work was funded by Royal Society/Wellcome Trust Sir Henry Dale Fellowship WT105677 (to VM). The authors declare no competing interests.

